# Magnetically Powered Microwheel Thrombolysis of Occlusive Thrombi in Zebrafish

**DOI:** 10.1101/2023.09.11.557256

**Authors:** M. Hao Hao Pontius, Chia-Jui Ku, Matthew Osmond, Dante Disharoon, Yang Liu, David W.M. Marr, Keith B. Neeves, Jordan A. Shavit

**Affiliations:** Department of Pediatrics, University of Michigan, Ann Arbor, MI; Department of Chemical and Biological Engineering, Colorado School of Mines, Golden, CO; Departments of Bioengineering and Pediatrics, University of Colorado, Denver | Anschutz Medical Campus, Aurora, CO; Department of Human Genetics, University of Michigan, Ann Arbor, MI

## Abstract

Tissue plasminogen activator (tPA) is the only FDA approved treatment for ischemic stroke but carries significant risks, including major hemorrhage. Additional options are needed, especially in small vessel thrombi which account for ∼25% of ischemic strokes. We have previously shown that tPA-functionalized colloidal microparticles can be assembled into microwheels (µwheels) and manipulated under the control of applied magnetic fields to enable rapid thrombolysis of fibrin gels in microfluidic models of thrombosis. Providing a living microfluidic analog, transparent zebrafish larvae have a highly conserved coagulation cascade that enables studies of hemostasis and thrombosis in the context of intact vasculature, clotting factors, and blood cells. Here we show that tPA-functionalized µwheels can perform rapid and targeted recanalization *in vivo*. This effect requires both tPA and µwheels, as minimal to no recanalization is achieved with tPA alone, µwheels alone, or tPA-functionalized microparticles in the absence of a magnetic field. We evaluated tPA-µwheels in CRISPR-generated plasminogen (*plg*) heterozygous and homozygous mutants and confirmed that tPA-µwheels are dose-dependent on plasminogen for lysis. We have found that magnetically powered µwheels as a targeted tPA delivery system are dramatically more efficient at plasmin-mediated thrombolysis than systemic delivery *in vivo*. Further development of this system in fish and mammalian models could enable a less invasive strategy for alleviating ischemia that is safer than directed thrombectomy or systemic infusion of tPA.

## Introduction

Ischemic stroke occurs in 700,000 people yearly in the United States, which often results in long-term disability(1, 2). Current treatments of ischemic stroke include systemic or catheter-mediated infusion of tissue plasminogen activator (tPA) which can lead to the dissolution of thrombi and subsequent recanalization of occluded vessels. tPA does this by converting plasminogen into plasmin which lyses fibrin(3). Systemic infusion however has a limited efficacy window and carries a significant risk of secondary hemorrhage at higher concentrations(1, 4).

Catheter-based thrombectomy is another option but is an invasive procedure with its own risks and cannot access small vessels(5, 6). As a result, additional treatment options are needed to minimize these risk factors and improve treatment outcomes, a goal especially important for small vessel thrombi which account for ∼25% of ischemic strokes(7).

The promise of micro-scale devices capable of medical intervention has led to the development of microbots that swim, crawl, and roll(8–11). Viscosity plays a dominant role in locomotion at small length scales(12), a factor that microorganisms overcome through physical adaptations, like rotating flagellum, that are difficult to artificially replicate and control(13, 14). In a particularly non-biomimetic approach, we have demonstrated a rapid and reversible microbot fabrication and powering method where µm-scale superparamagnetic beads assemble upon application of a relatively weak (1-5 mT) rotating magnetic field(15). In this, superparamagnetic beads experience strong attractive interactions, bringing them together to assemble into disc-like shapes we call µwheels. These µwheels roll rapidly (100’s µm/sec) and can be immediately redirected with a simple alteration in the magnetic field orientation resulting in speed and heading changes. Using different time varying magnetic fields, µwheel size distributions can be created and specific modes specializing in various tasks designed(16). Four unique µwheel modes corresponding to specific needs are discussed here: rolling mode, for optimal mass flux; switchback mode, for steep incline traversal; flipping mode, for deposition of small microbots across a large area; and corkscrew mode, to support burrowing in porous structures.

We have previously shown that µwheels functionalized with tPA enable rapid thrombolysis of fibrin gels and platelet-rich thrombi in vitro by allowing for higher local concentrations of tPA and mechanical burrowing into a thrombus(15, 17). Corkscrew mode demonstrated superior penetration into fibrin gels compared to rolling mode(15). However, while we were able to achieve local tPA concentrations that were 3 orders-of-magnitude higher than those reached by diffusive delivery of tPA, this resulted in only a 1 order-of-magnitude increase in fibrinolysis speed. Based on these results we showed that there is a biochemical speed limit of fibrinolysis due plasminogen depletion at high tPA concentrations(18). We have yet to show the efficacy of µwheels for thrombolysis and whether the same plasminogen depletion occurs in vivo.

Zebrafish (*Danio rerio*) is a vertebrate with significant homology to mammalian coagulation systems(19). Development is rapid, external, and embryos and larvae are transparent in the early stages of life, making zebrafish essentially a living microfluidic model(20). This allows for studies using an optical microscope in vivo with intact vasculature, coagulation factors, and blood cells in the context of normal and disrupted hemostasis(21). Here, we show that tPA-functionalized µwheels infused into zebrafish larvae can achieve semi-targeted recanalization *in vivo*, with the corkscrew mode of translation yielding the fastest recanalization times.

Thrombolysis is achieved at lower overall concentrations compared to tPA infusion and is dependent on plasminogen.

## Results

### µWheels facilitate tPA-dependent recanalization *in vivo*

We previously demonstrated that only µwheels functionalized with tPA can achieve lysis of fibrin clots *in vitro*(17). To test this *in vivo*, we first infused fluorescently tagged microparticles into circulation at 5 days post fertilization (5 dpf) through the retro-orbital plexus (Fig. 1E). These were observed for 1 hour in circulation with no adverse effects nor any accumulation in any particular location observed. We have previously shown the ability to produce occlusive thrombi using laser-mediated endothelial injury in 3-5 dpf zebrafish larvae^13^, as well as the ability to lyse spontaneous thrombi using human tPA(22). We performed injury 1 hour after microparticle infusion and found accumulation at the upstream edge of the occlusive thrombus (Fig. 1F).

**Figure 1.**
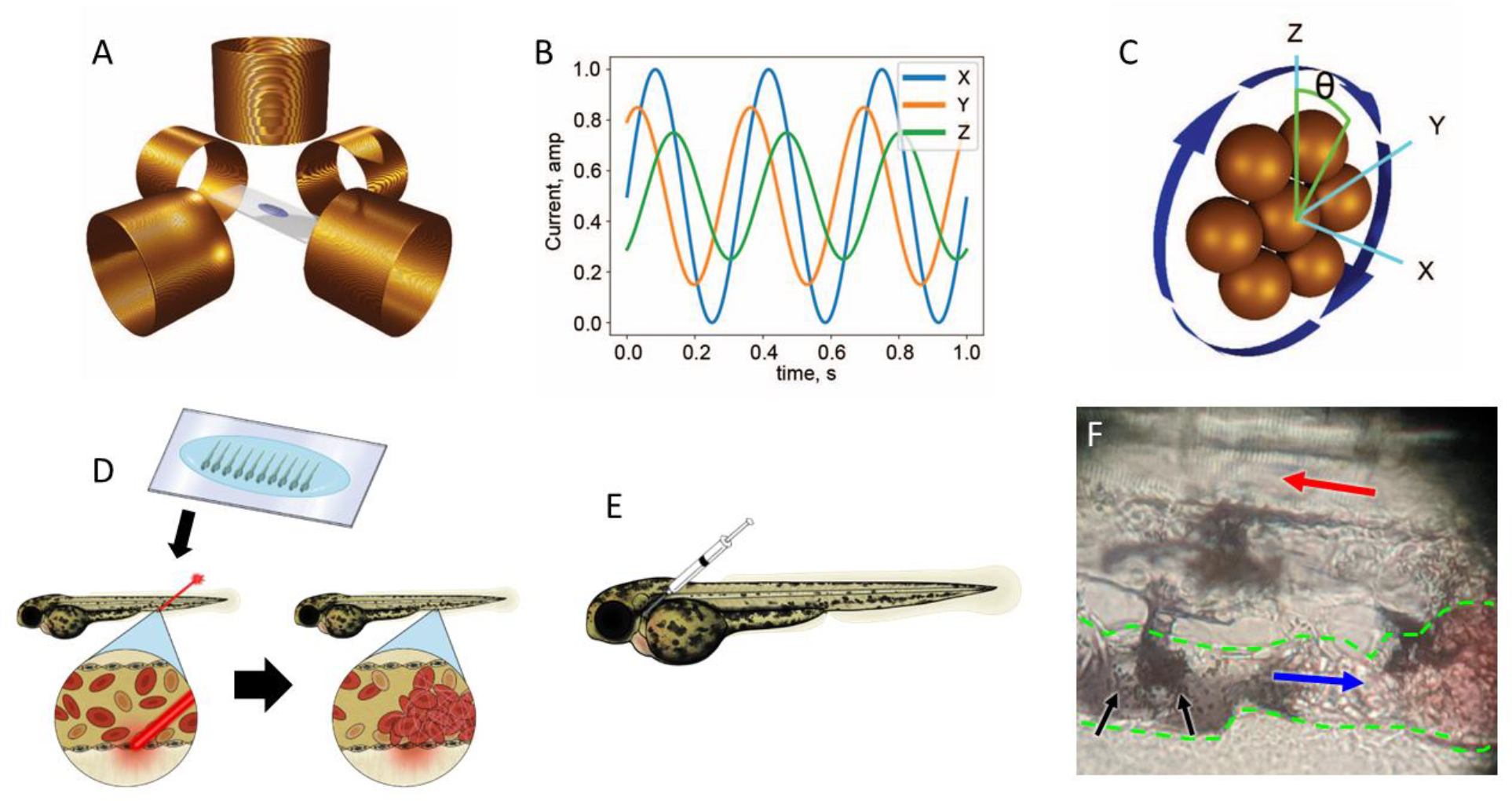
Generation of µwheels. (A) zebrafish larvae mounted in low melting point agarose and infused with microparticles were placed in the center of 5 solenoid coils. (B) A laptop running muControl software connected to a function generator and amplifier sends alternating currents to the x, y, and z coils. (C) The rotating magnetic field (blue arrows) induced by the coils causes the assembly and translation of the µwheels with steering controlled by the camber angle (θ) of the field relative to the z-axis. (D) 3 dpf zebrafish larvae were mounted in low melting point agarose on glass cover slips and a pulsed dye laser was used to injure the endothelium of the PCV, resulting in thrombosis. (E) After endothelial injury, tPA and/or microparticles were infused via the retro-orbital space or PCV using pulled capillary micropipettes. (F) Photograph of µwheels at the site of a clot. Red and blue arrows indicate the direction of blood flow in the arterial and venous systems, respectively. Black arrows indicate the µwheels at the upstream edge of an occlusive thrombus, outlined with green dashes.

These studies demonstrated that zebrafish larvae tolerate circulation of the beads and that they can be easily visualized.

While 1 ng of tPA infused into individual larvae is approximately equal to the dosing in patients (0.9 mg/kg) we tested a range of doses, from 1-40 ng/larvae, for recanalization. Only high concentrations of tPA, greater than 10 ng/larvae, were able to achieve recanalization within 4 hours and without significant toxicity (Fig. 2). Greater concentrations resulted in toxicity evidenced by gross morphologic changes. These studies suggest that, to achieve efficient lysis with minimal toxicity, a high local concentration of tPA is needed.

**Figure 2.**
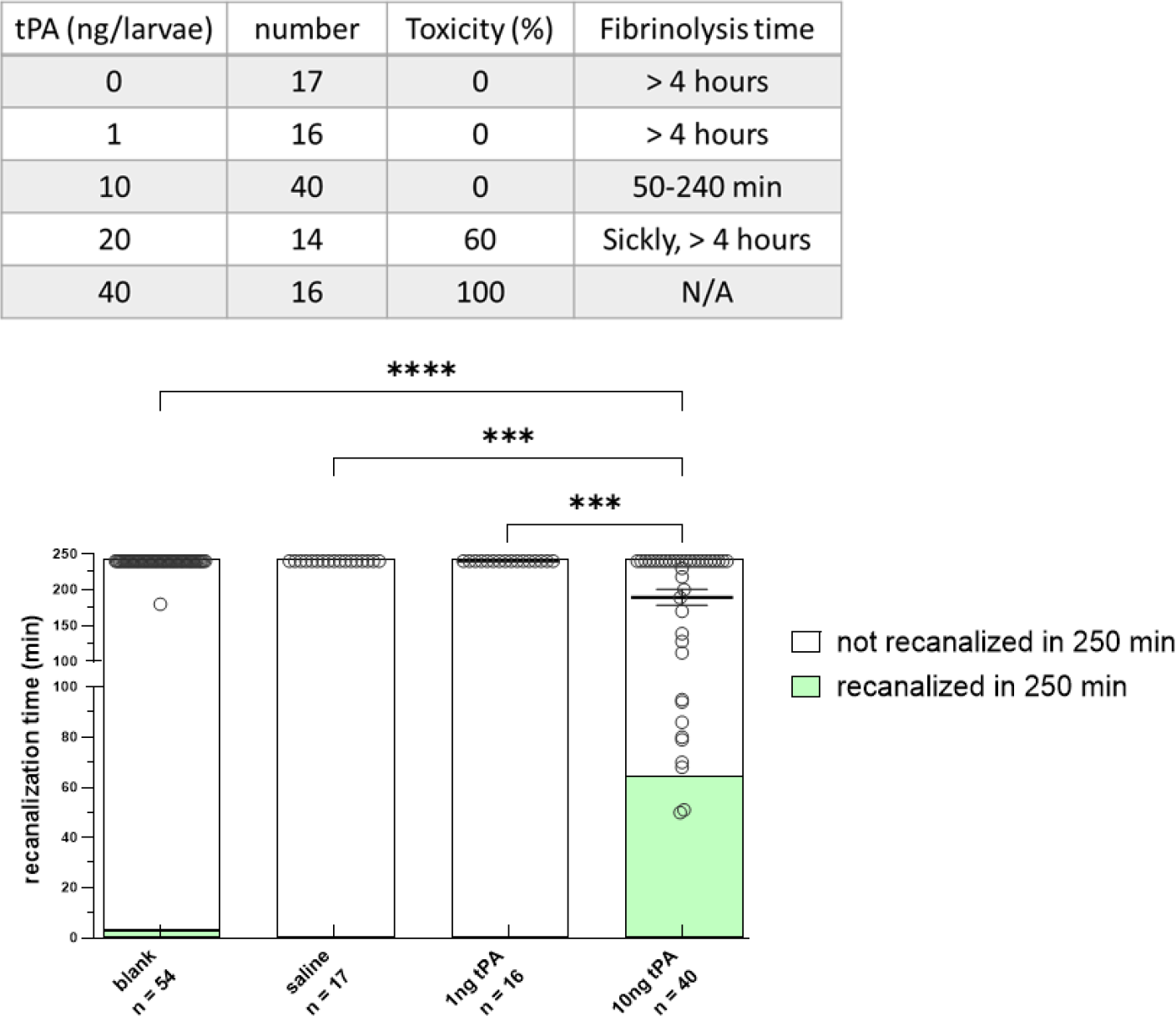
Time to fibrinolysis after infusion of tPA. The PCV of 3 dpf larvae was injured, producing a completely occluding thrombus. This was followed by either infusion with recombinant human tPA or microparticles functionalized with tPA, and then observation for up to 4 hours. Only high concentrations of tPA at 10 ng/larvae (∼5-10x the amount used for human therapy) were able to achieve recanalization in a fraction of larvae within 4 hours and without significant toxicity. Lower concentrations of tPA and microparticles functionalized with tPA but without the magnetic field, did not recanalize except for one in the latter group. Bars represent mean recanalization time. Bar graphs indicate the percentage of clots that recanalized. (**p < 0.005, ***p < 0.0005, ****p < 0.0001 by Kruskal-Wallis testing).

To test the ability of µwheels to recanalize the vessel, we infused microparticles functionalized with recombinant human tPA or Atto-488 fluorescent dye, followed by endothelial injury 1 hour after infusion, and then application of the magnetic field to produce µwheels. All of the larvae with tPA functionalized µwheels recanalized within 15 minutes, while control µwheels did not recanalize within 30 minutes (Fig. 3, Mov. S1A-D). We infused tPA-functionalized microparticles without application of a magnetic field (Fig. 1E) and found no evidence of significant recanalization within 4 hours. Next, we infused non-functionalized particles followed by application of a magnetic field using corkscrew motion. We were only able to apply the magnetic field for 30 minutes given the heat generated by the coils, but there was no lysis in the absence of tPA. Taken together, these data show that only tPA-functionalized µwheels are capable of thrombolysis.

**Figure 3.**
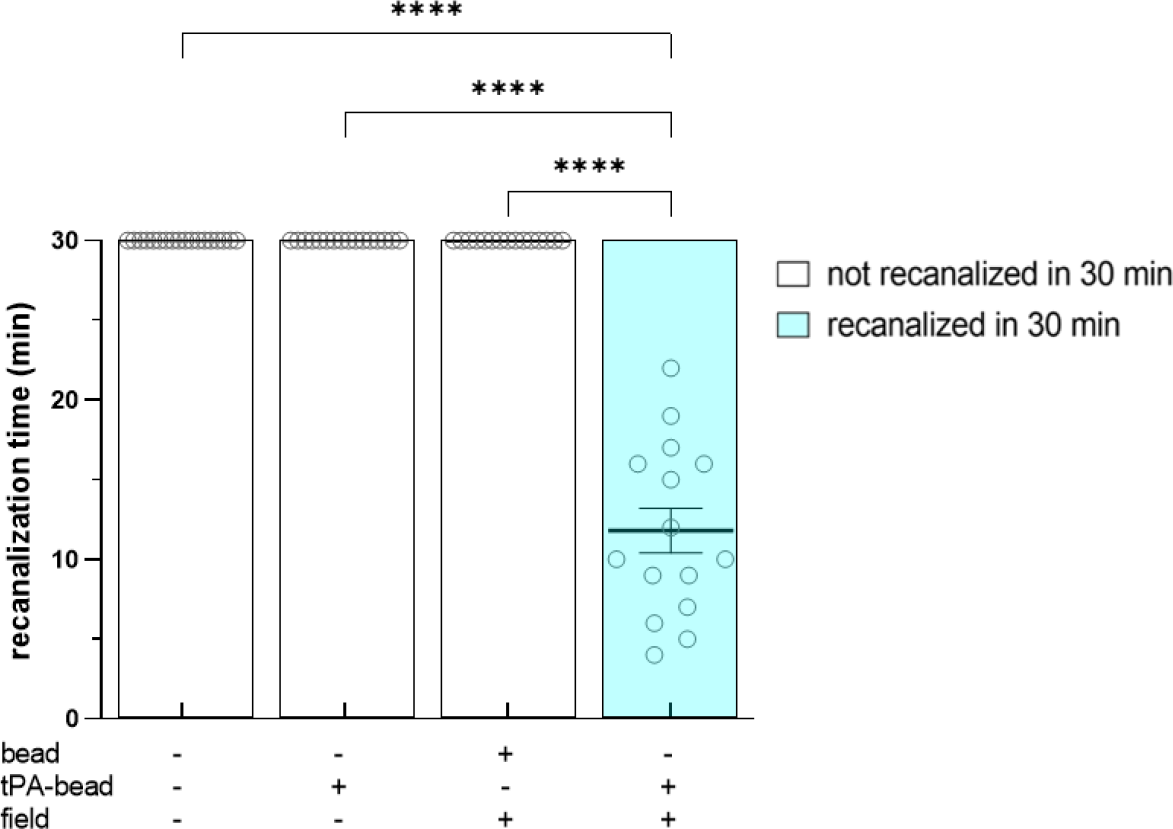
tPA-functionalized µwheels are necessary for recanalization. The PCV of 3 dpf larvae was injured, producing completely occlusive thrombi, followed by the infusion of unfunctionalized microparticles, microparticles functionalized with tPA, or uninfused controls, and in some cases the application of a magnetic field to produce µwheels. The field was applied in rolling mode either until recanalization was observed or a maximum of 30 minutes. Beyond that point the coils became too hot and damaged the larvae. Larvae infused with tPA-functionalized µwheels recanalized within 11.8 ± 5.4 minutes while those infused with non-functionalized µwheels, tPA functionalized particles without the application of a magnetic field, and uninfused larvae continued to be occluded after 30 minutes (p < 0.0001 by Mann-Whitney U testing). Bars represent mean recanalization time. Bar graphs indicate the percentage of clots that recanalized.

### Corkscrew motion of µwheels is most effective for recanalization

The corkscrew motion has been previously shown to be the most effective at lysis *in vitro*(15). To examine if this is also the case *in vivo*, we performed a comparison to additional modes: switchback, flipping, and rolling movements. We compared all of these using functionalized µwheels and found that corkscrew motion was most effective at recanalizing occlusive thrombi, in a median time of under 10 minutes. The other modes were also able to recanalize occlusive thrombi within 30 minutes, but with a median of ∼15 minutes or higher (Fig. 4).

**Figure 4.**
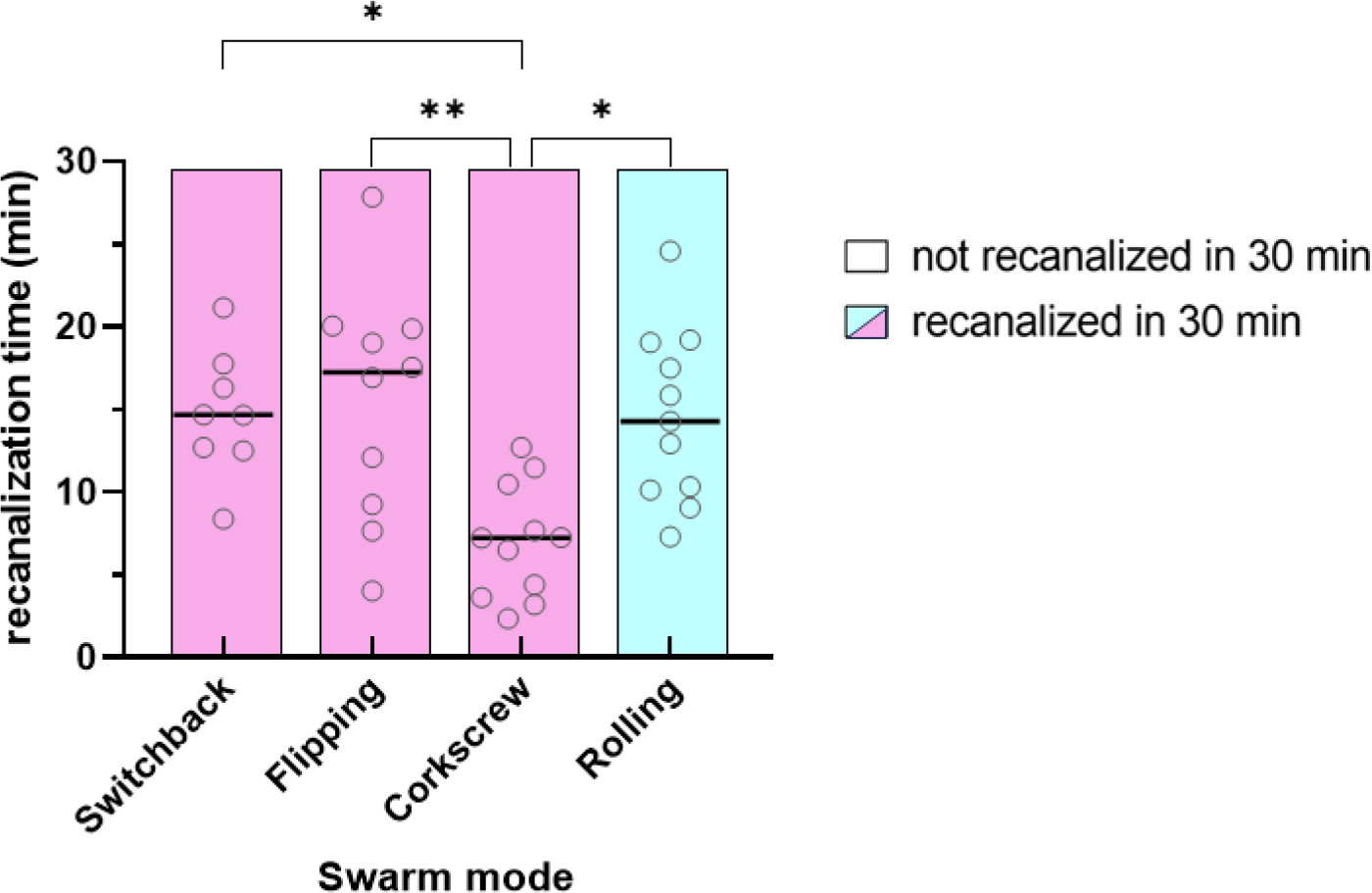
Effect of motion type on recanalization time. µwheels under “switchback”, “flipping” or “rolling” modes of movement all recanalize within 30 minutes, while µwheels under “corkscrew” movement recanalize within 15 minutes (*p < 0.05, **p < 0.01 by Kruskal-Wallis testing). Bars represent mean recanalization time. Bar graphs indicate the percentage of clots that recanalized, blue indicating “rolling” motion and pink indicating other modes.

### µWheel-mediated recanalization of occlusive thrombi is dependent on plasminogen

tPA is well known to mediate its effects by activating plasminogen to plasmin. Although the plasminogen (*plg)* gene is single copy and conserved in the zebrafish genome, it has not been previously shown whether it has the same function. Microparticles were infused into larvae generated from *plg*^*+/-*^ incrosses after endothelial injury, followed by application of the magnetic field. µWheels were able to recanalize occlusive thrombi in the majority of wild-type and heterozygous mutants within 30 minutes, while nearly all of the homozygous mutants were unable to do so (Fig. 5). There was a small significant decrease in the percentage of heterozygous mutants that were able to recanalize compared to wild-type. These data show that fibrinolysis is plasminogen-limited, at least at high local tPA concentrations(18).

**Figure 5.**
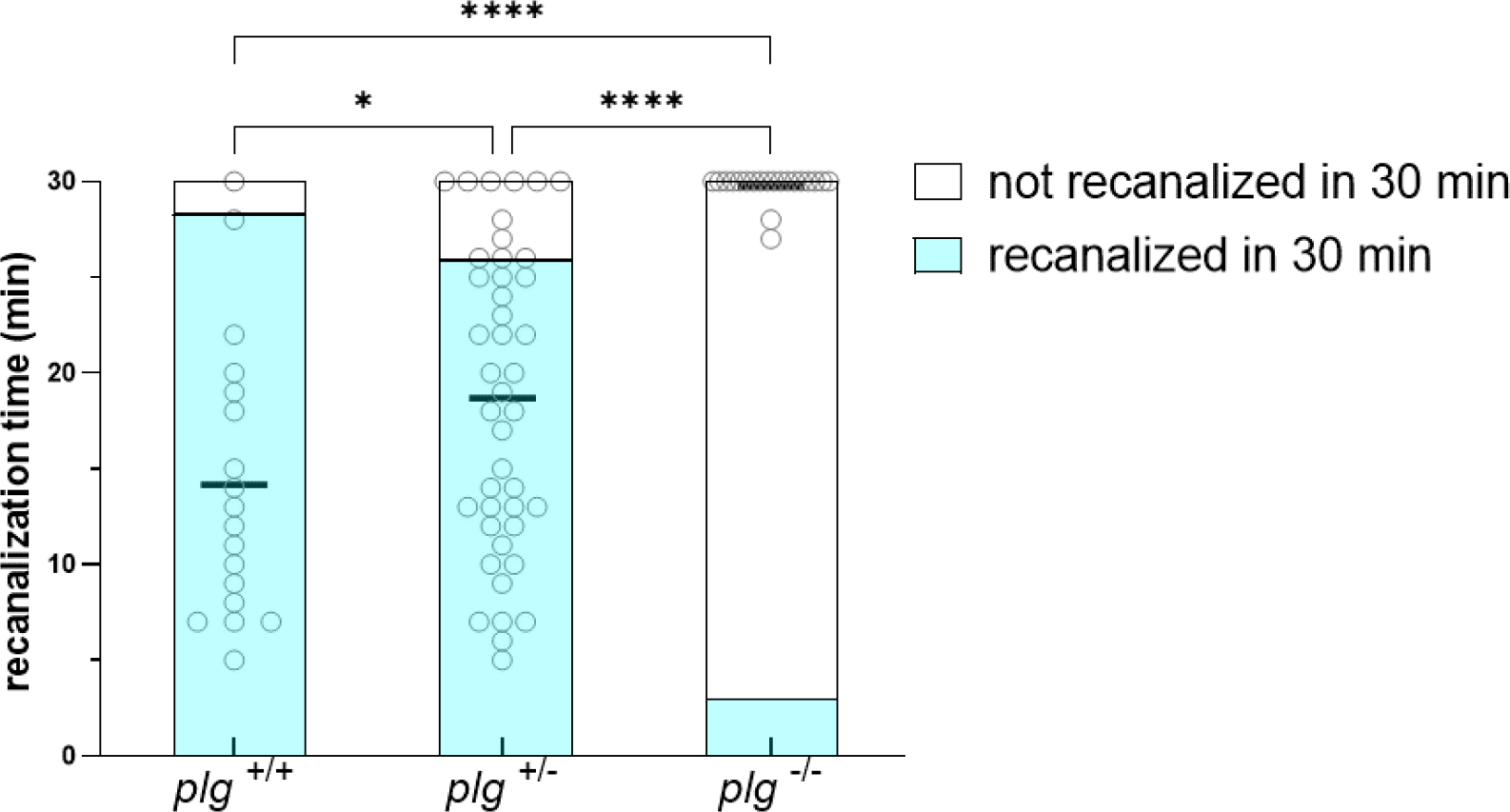
µwheel-mediated recanalization of occlusive thrombi is dependent on plasminogen. CRISPR-mediated genome editing was used to produce a 23 base pair deletion in the *plg* gene. In the majority of wild-type and heterozygous mutants, µwheels were able to recanalize an occlusive thrombus within 30 minutes. Conversely, the vast majority of homozygous mutants were unable to do so. Interestingly there was a small, but statistically significant decrease in the percentage of heterozygotes that were able to recanalize, when compared to wild-type siblings. Observations were performed prior to genotyping, and thus were blinded. Bars represent mean recanalization time. Bar graphs indicate the percentage of clots that recanalized under “rolling” mode. (*p < 0.05, ****p < 0.0001 by Mann-Whitney *U* testing).

## Discussion

Within a limited timeframe of 3-4.5 hours after onset(23), systemic or localized administration of tPA is a current treatment for ischemic stroke which works through the conversion of plasminogen into plasmin, which then effects fibrinolysis. While localized adminsitration is limited to large vessels, systemic delivery has a risk of widespread plasminogen activation, which can result in hemorrhage at high concentrations. A method therefore of delivering high concentrations of tPA directly to thrombi while maintaining low systemic concentrations of tPA could be safer and more effective. In our model, magnetically powered µwheels as a targeted tPA delivery system are dramatically more efficient at plasmin-mediated thrombolysis than systemic administration. In humans, this approach would enable a less invasive strategy for alleviating ischemic stroke that is safer than directed thrombectomy or systemic infusion of tPA. We have previously shown that µwheels are effective in uniform *in vitro* fibrin gels(15). Here we demonstrate that they can also facilitate thrombolysis in an *in vivo* system in the context of intact vasculature, coagulation factors, and blood cells. Surprisingly, most of the features are similar. In both systems, tPA-functionalized µwheels are required, and tPA-microparticles or non-functionalized µwheels are ineffective. Corkscrew motion is also the most efficacious both *in vitro* and *in vivo*. Notably, the total concentration of tPA delivered is far lower than the amount needed for lysis with systemic therapy (at levels analogous to human infusions) in this model.

While the rotating magnetic field controls the assembly and rolling direction of µwheels, we determined blood flow and accumulation of µwheels in low flow regions proximal to the thrombus sufficient to drive thrombolysis. In systems with higher flow rates, magnetophoretic forces can be added to direct µwheels/microparticles to the thrombus with electromagnets(24) or with a rotating permanent magnet when significantly higher fields are desired(25).

Using genome editing, we produced a model of plasminogen deficiency. As expected, we were able to confirm that the functionalized µwheel effects are due to the production of plasmin.

Homozygous *plg* mutants had almost no recanalization, verifying conservation of the fibrinolytic system in zebrafish and the ability of human tPA to activate fish plasminogen. Interestingly, we found that there was a dose dependent response, as heterozygotes had a mean recanalization time that was significantly longer than wild-type siblings. This is consistent with plasminogen being the rate-limiting component in local fibrinolysis, as we have previously shown(18). When there is excess tPA, the lytic rate is dictated by available plasminogen. This is consistent with clinical practice and data in which plasminogen levels are correlated with tPA therapeutic success(26, 27).

In summary, we show that zebrafish are an effective transitional model between *in vitro* systems and more complex mammalian models. They are like living microfluidic systems, whose transparency and vasculature enable simple imaging. The results demonstrate a high degree of concordance with microfluidic models, promising a future rapid *in vitro* to *in vivo* pipeline.

### Materials and Methods Animal Care

Zebrafish were maintained according to protocols approved by the University of Michigan Animal Care and Use Committee. Wild-type fish were a hybrid line generated from crosses of AB and TL zebrafish acquired from the Zebrafish International Resource Center. Tricaine (Western Chemical) was used for anesthesia and rapid chilling in an ice water bath for euthanasia.

### Functionalization of microparticles

Recombinant tPA was biotinylated using N-hydroxysuccinimide (NHS) activated biotin by incubation on ice for 3 hours. Excess biotin was removed using a desalting column, and biotinylated tPA stored in 20 µL aliquots at -80°C. 5 µL of 1 µm streptavidin-conjugated iron oxide beads (ThermoFisher Dynabeads MyOne Streptavidin T1) were mixed with biotinylated tPA, incubated at 4°C overnight, washed three times to remove excess tPA, and resuspended in 15 µL 2% bovine serum albumin (BSA) in saline. Atto 488-biotin was similarly conjugated to Dynabeads without tPA functionalization and used as a control.

### Laser-mediated endothelial injury assay

Laser-mediated endothelial injury was performed on 3 dpf zebrafish larvae(20). Larvae were anesthetized in tricaine and mounted in 0.8% low melting point agarose on glass cover slips. The agarose around the head of the larvae was removed and replaced with water. A laser (MicroPoint Pulsed Laser System, Andor Technology, Belfast, Northern Ireland) was used to injure the endothelium of the posterior cardinal vein (PCV) 5 somites caudal to the anal pore using 99 pulses.

### Infusion of tPA and microparticles

After endothelial injury, 1-40 ng tPA or 3 nL microparticles were infused via retro-orbital vessels using pulled capillary micropipettes(20) (Figs. 1D and E). Standard tPA dosage was 0.9 ng/mg (6 dpf larvae have been shown to weigh ∼1 mg(28), and 3 dpf is estimated to be 0.5-1 mg), corresponding to clinical dosages of 0.9 mg/kg(3). Functionalized µwheels carried ∼0.7-2.1 pg tPA/µwheel(18).

### Generation of µwheels

After introduction of the microparticles into the zebrafish larvae, µwheels were assembled using a rotating magnetic field generated with a custom apparatus(29). This apparatus consists of 5 coils arranged with two opposing pairs in the x and y plane and a fifth coil in the z-direction above the zebrafish sample. A set of sinusoidal currents are produced by an analog output card (NI-9263, National Instruments, Austin, TX), run through an amplifier (EP2000, Behringer, Willich, Germany), and sent to each pair of coils producing a rotating magnetic field. The circular, rotating magnetic field is positioned normal to the rolling surface and the direction of the µwheels manipulated by varying the ratio of current supplied to the individual coils. The magnetic field strength and frequency are controlled with custom software(30). Previously, four different magnetic field modes were investigated for controlling µwheel swarms, including switchback, flipping, corkscrew, and rolling(16). Here we used these modes to investigate their effect on the time to recanalization. Switchback mode results from switching the heading angle of the wheel back and forth which provides better movement over inclined surfaces. Flipping results from changing the camber angle back and forth and provides a way for wheels to prevent assembly into larger wheels. Corkscrew uses a synchronized change in heading angle and camber together that results in enhanced penetration. Rolling is our standard mode in which the heading is fixed manually to control directed movement through complex geometries (Fig. 1)(16, 29).

### Measurement of recanalization

Time to recanalization was observed optically for up to 30 minutes and defined by the resumption of passage of blood cells beyond the occlusive thrombus and into the distal PCV. Longer times were not feasible due to coil heat generation. All observations were collected by an observer blinded to condition or genotype.

### CRISPR/Cas9 mediated genome editing

We used CRISPR/Cas9 mediated genome editing system to produce a 23 base pair deletion in the plasminogen (*plg*) gene, as previously described(31). Briefly, single guide RNAs (sgRNAs) were identified using ZiFiT Targeter(32), and sequence GGGAGTACTGCAATATTGAG in exon 12 was selected. DNA templates were produced using pDR274(33), and sgRNAs transcribed. sgRNAs and Cas9 mRNA were co-injected into one-cell stage zebrafish embryos, at concentrations of 12.5 and 300 ng/µL, respectively. F0 offspring were raised to adulthood and crossed with wild-type zebrafish to verify F1 germline transmission. Heterozygotes were incrossed to produce homozygous mutants.

### Genotyping of offspring

Genomic DNA of zebrafish larvae was isolated after incubation in lysis buffer (10 mM Tris-Cl, pH 8.0; 2 mM EDTA, 2% Triton X-100, and 100 µg/mL proteinase K) at 55°C for 2 hours followed by proteinase K inactivation at 95°C for 5 minutes. Mutations were detected by PCR using primers 5’-AGATCTAAGGAGAAACCTGT-3’ and 5’-CTTTTTCTGAGGGAGCAGAT-3’.

### Statistical analysis

Statistical analysis was performed with Mann-Whitney *U* or Kruskal-Wallis testing. Significance testing was performed using Prism (GraphPad software, California).

## Acknowledgements

This work was supported by National Institutes of Health (NIH) grants R01 NS102465 to D.W.M.M and K.B.N. and R35 HL150784 to J.A.S. J.A.S. is the Henry and Mala Dorfman Family Professor of Pediatric Hematology/Oncology.

## Author Contributions

M.H.H.P. designed and performed research, analyzed data, and wrote the manuscript, C.J.K. designed and performed research and analyzed data, M.O. and D.D. designed research and analyzed data, Y.L. designed and performed research, D.W.M.M., K.B.N., and J.A.S. designed and supervised research and edited the manuscript.

---

**Movie S1. µwheels in circulation** (A) Laser-mediated endothelial injury was performed in the PCV of 3 dpf larva, producing an occlusive thrombus. Circulation is flowing through collateral vessels. (B) Microparticles were infused through retro-orbital vessels and are shown in circulation without the application of a magnetic field. (C) µwheels were formed after application of the magnetic field and can be seen circulating in large clusters. (D) An occlusive thrombus was produced in the PCV as in (A). tPA-functionalized microparticles were infused followed by application of the magnetic field to produce µwheels, resulting in thrombolysis and vessel recanalization.

